# A Novel COE-D8-Fosfomycin Conjugate Effectively Against First-line Antibiotic Resistant *Uropathogenic Escherichia coli*

**DOI:** 10.1101/2025.01.19.633753

**Authors:** Lin Ruan, Cun Fan, Jingjie Chen, Yingxia Wang, Yuji Ren, Wenli Yuan, Xiaozhe Xiong, Chunming Guo

## Abstract

Recurrent urinary tract infections (UTIs) are increasingly associated with antibiotic resistance, posing significant clinical challenges. This study evaluates the antimicrobial efficacy of the membrane-intercalating conjugated oligoelectrolyte COE-D8 against uropathogenic Escherichia coli (UPEC). Among 93 clinical UPEC isolates, over 50% demonstrated resistance to first-line antibiotics, with compound sulfamethoxazole showing the highest resistance (75% at a minimum inhibitory concentration (MIC) >512 μg/mL). COE-D8 significantly reduced resistance rates, achieving less than 50% at 32 μg/mL and below 5% at 512 μg/mL, while demonstrating at least fourfold faster bacterial elimination than standard antibiotics.

Mechanistic studies revealed that COE-D8 disrupts bacterial membranes, inducing rapid lysis. However, its cytotoxicity in mammalian cells at 2 μg/mL limits clinical application. To address this, a combination therapy with fosfomycin was explored, significantly reducing COE-D8 toxicity by at least fourfold in vitro. In mouse models, fosfomycin co-administration mitigated COE-D8-induced toxicity and prevented mortality at higher doses without compromising antimicrobial efficacy.

These findings highlight the potential of COE-D8, particularly in combination with fosfomycin, as a promising strategy to combat antibiotic-resistant UPEC while addressing toxicity challenges.

## 1. Introduction

Urinary tract infections (UTIs) are common and significantly impact the quality of life, particularly for women, due to the anatomical structure of the female lower urinary tract and its close proximity to the reproductive organs.^1^The incidence of UTIs ranks second only to respiratory and digestive tract infections across all populations.^2 3^Even after antibiotic treatment, 30 to 50% of patients continue to suffer from recurrent attacks^4^. *Escherichia coli* (E. coli), particularly uropathogenic E. coli (UPEC), is the primary “culprit” behind urinary tract infections. UPEC initially infects bladder epithelial cells (BECs) via the unique spindle vesicles present in these cells, which play a crucial role in exocytosis and contribute to both physiological and immune functions.^5^To evade exocytosis-mediated clearance, pathogenic E. coli escapes from the spindle vesicles, enters the cytoplasm, and replicates extensively to form an intracellular bacterial community containing over 10,000 bacteria. This enables pathogenic E. coli to successfully colonize the bladder epithelium and initiate infection.^6–12^

In clinical practice, fosfomycin has been approved for treating uncomplicated UTIs. It is administered as a single oral dose of 3g, offering good tolerability and achieving high concentrations in the urine.^13–16^Fosfomycin has a unique chemical structure and does not exhibit cross-resistance with other drugs, which has garnered increasing interest in its expanding role in the treatment of multidrug-resistant UTIs ^17–19^ and as a therapeutic option for UPEC-associated UTIs.^20,21^

Despite the widespread use of fosfomycin and other antibiotics in the treatment of UTIs, resistance among UPEC strains has grown,^22^making it a significant challenge. This resistance has extended to nearly all classes of antibiotics, including recently developed drugs like oxacillin, flucloxacillin and dicloxacillin.^23^This resistance poses significant challenges for anti-infection therapy, as it hinders antibiotics from penetrating bacterial communities, further increasing bacterial resistance. As a result, it is increasingly difficult for antibiotics to penetrate bacterial communities, further exacerbating resistance. Despite extensive efforts to develop new antibiotics,^14^ no new class of antibiotics has been approved for the treatment of Gram-negative bacterial (GNB) infections in over 50 years.^24^Antimicrobial peptides (AMPs) are believe to kill bacteria by destabilizing their membranes or cell walls through multiple complementary mechanisms, which help mitigate the development of resistance. ^25–27^ However, their toxicity often deters widespread use, making commercialization difficult.^28,29^Therefore, the combination therapy involving antimicrobial compounds and clinical antibiotics has emerged as a promising strategy to address both toxicity and drug resistance.

In our previous work, we reported that Conjugated oligoelectrolytes (COEs) represent a new class of antibiotics that exhibit superior efficacy against *MRSA* compared to vancomycin.^30^Specifically, COEs consist of a hydrophobic, linear conjugated backbone (such as oligo-phenylenevinylene) with terminal ionic groups. Their molecular structure closely resembles the distribution of charged and hydrophobic domains in lipid bilayers, enabling COEs to spontaneously intercalate and partition into cellular membranes.^31^ The length of a COE influences its effects on lipid bilayers after intercalation.^32^As shown in Scheme S1^30^, COE-D8 is a typical and potent antimicrobial molecule, with its antimicrobial activity likely resulting from membrane disruption caused by a dimensional mismatch between the COE length and the lipid bilayer thickness.^33^Consequently, COE-D8, can exhibit antimicrobial properties against clinical UPEC by disrupting the bacterial cell membrane.

In this study, we demonstrate the unexpected synergistic effect of combining the “disruptor” COE-D8 with the “stabilizer” fosfomycin, which results in reduced cytotoxicity and enhanced antimicrobial efficacy, providing a promising foundation for subsequent therapeutic interventions.

## 2. Materials and methods

### 2.1 Materials, strains and instruments

The COE-D8 used in this study was synthesized according to previous literature protocols^30,34^. The following drugs were purchased from MCE: Colistin, Polymyxin B, Ampicillin, Kanamycin, Fosfomycin, Cephalothin, Compound Sulfamethoxazole, Levofloxacin, and Nitrofurantoin. Experimental mice were obtained from the Animal Center of Yunnan University (Ethics approval number: CHSRE20220307). Ninety-three strains of multidrug-resistant *Uropathogenic Escherichia coli* (MDR-UPEC) were isolated from urine culture plates of patients in the Department of Urology, The First Affiliated Hospital of Kunming Medical University. Mueller Hinton Broth (MHB, Solarbio), Luria-Bertani(LB, Solarbio) and (1×) Phosphate Buffered Saline (PBS, pH = 7.2) were prepared and autoclaved before use. The 3T3 cell line, purchased from Pricella (Wuhan, China), was used as a mammalian cell model in the *in vitro* cytotoxicity assays. Hematoxylin and eosin (HE) tissue sections were prepared using a paraffin embedding machine (MEDITE TES99), followed by visualization with a Slide Scanning System (Shenzhen Shengqiang Technology Co, Ltd., SQS40P/SQS40PRO).

### 2.2 Bacteria isolation, culture and reservation

Upon obtaining 93 culture plates from the hospital, half of the bacteria from each plate were directly scraped into a 50% glycerol solution, designated as the “O” series (O stands for “original”), e.g., 1O, 2O, and 3O. The remaining bacteria were scraped into 20 mL of LB medium and incubated overnight at 37°C with shaking at 220 rpm. After 24 hours, the culture was mixed with 50% glycerol and stored at −80°C, labelled as the “L” series. Subsequently, the frozen “L” series bacterial cultures were retrieved from the −80°C freezer and a small amount was transferred into 5 mL of LB/MHB medium using a sterile loop. This was then incubated overnight at 37°C with shaking at 220 rpm. After another 24 hours, a portion of the overnight culture was diluted 1:10 or 1:5 and transferred into 5 mL of fresh LB/MHB medium for subculture. The bacteria were allowed to reach the logarithmic phase, followed by a tenfold serial dilution. A 10^3 dilution was plated onto a 90 mm LB agar plate and incubated overnight at 37°C. The next day, three distinct colonies from each plate were selected and stored, followed by a repeated 24-hour growth cycle for further analysis.

### 2.3 Method

#### 2.3.1 Minimum Inhibitory Concentration (MIC) determination

MIC values are determined using a broth micro dilution method. Bacteria cells are grown overnight at 37 °C in MHB to a mid-log phase (OD_600_ between 0.4-0.5 for each organism) and diluted in MHB to 10^5^ CFU/mL. The COE-D8 and different antibiotics is dissolved in corresponding solvent to a stock concentration of 10 mg/mL. The antibiotics listed in Table S1 are dissolved and prepared to stock concentrations according to CLSI guidelines. 50 µl of the 1-5×10^5^ CFU/mL bacterial cultures (final concentration) is aliquoted into 96-well microtiter plates and mixed with 50 µl of two-fold dilutions of the COE-D8 or antibiotics and incubated for 16-18 h at 37°C with shaking at 200 rpm. Growth inhibition is determined by measuring the optical density at 600 nm (OD_600_) of each well using a TECAN M200 microplate reader; the lowest polymer concentration at which exhibited no bacterial growth is defined as the MIC.

#### 2.3.2 Mammalian Cell Biocompatibility test via MTT cell proliferation assay

The mammalian cell biocompatibility test is done according to the published protocol using 3T3 cells^35^. In a 96-well plate, 3T3 cells are co-cultured for 24 hrs at 37°C with COE-D8 (from 1 μg/mL to 512 μg/mL) at initial cell density of 1×10^5^ cells per well. At the end of the incubation period, the culture medium is removed, each well is washed with PBS followed by the addition of MTT solution, and the plate is incubated for 4 h at 37°C. The MTT medium is then removed, 100 μL of DMSO is added to each well, the plate is shaken at 100 rpm for 15 mins and the absorbance at 570 nm is measured with plate reader (TECAN M200, Switzerland).

#### 2.3.3 Time Killing assay

Bacteria cells are grown, diluted, and aliquoted into 96 well plates as described for the MIC assay, and then mixed with 50 µl volume of medium containing 4 × COE-D8 and antibiotics for NO. strain 63,73,76 and 80 or containing 1 × MIC COE-D8 and Fosfomycin combination for NO. strain 73 and 76. The plates are sealed and incubated at 37°C with shaking at 200 rpm.

At 0, 0.5, 1, 2, 3, 4, 5 and 6h post-inoculation, each well is thoroughly mixed with a multi-channel pipette and 20 μl of sample is removed, serially diluted in sterile phosphate buffered saline (PBS), plated on LB agar plates, and incubated at 37°C for 18 hrs. Colonies are counted to determine the CFU/mL at each time point.

#### 2.3.4 Evolution of COE-D8 resistance

The evolution experiment for spontaneous COE-D8 resistance was performed through sequential passaging as described elsewhere^36^. UPEC strains (#63, #74, #76, #80) grown in MHB medium were used. The initial inoculum for the first transfer was 10^7 cells/mL in a 24-well plate, with varying concentrations of COE-D8, and a total volume of 1 mL. Growth was monitored at 24-hour intervals. Daily transfers were performed, with the inoculum (100-fold dilution) taken from cultures where the antibiotic concentration allowed growth to an OD_600_ of at least 0.2, followed by exposure to higher concentrations of the compound. The experiment for strains #63 and #76 lasted 33 days, for strain #74 the experiment ended at 19 days, and for strain #80, the experiment continued until day 28. Isolates from the final transfer of the four strains were stored as glycerol stocks at −80°C for further studies.

#### 2.3.5 Outer-membrane permeability assay

1-N-phenylnaphthylamine (NPN) assay was conducted on Gram-negative bacteria UPEC #73 and #76 to determine the ability of COE-D8 to permeabilize outer-membranes^9^. Bacteria overnight cultures were prepared by inoculating a few colonies in MHB. A pre-culture of bacteria was prepared by diluting overnight culture to an OD_600_ of 0.01 in fresh MHB; this was incubated with shaking at 37°C, 200 rpm to mid-log phase (OD_600_= 0.5). Bacterial cells were then collected by centrifugation at 3,000 X g for 10 min, washed twice with 5 mM HEPES buffer and resuspended to a final concentration of 10^7^ CFU/ml in 5 mM HEPES buffer. 50 µl of 40 µM NPN solution was added to 100 µl of bacteria suspension in a 96-well black plate (Corning Costar). 50 µl of poly(beta-peptide) in HEPES buffer solution (at desired concentration) was added, and fluorescent signal was immediately measured (TECAN, infinite F200) at an excitation wavelength of 350 nm and an emission wavelength of 420 nm. The following negative controls were also recorded: 100 µl of bacteria + 100 µl of HEPES buffer; 50 µl of 40 µM NPN + 150 µl of HEPES buffer; 100 µl of bacteria + 50 µl of HEPES buffer + 50 µl of 40 µM NPN.

#### 2.3.6 Animal Cytotoxicity assay

Seven-week-old female ICR mice were used as experimental subjects, all of which were purchased from the Animal Center of Yunnan University (Ethical Code: CHSRE2022030). Prior to intraperitoneal injection of the drugs, the mice were weighed to establish the baseline at 0 hours. Subsequently, their weight was measured every 24 hours for a total of 7 consecutive days. The experimental groups were as follows: (1) 25 mg/kg COE-D8 alone, (2) 50 mg/kg Fosfomycin alone, and (3) a combination of 25 mg/kg COE-D8 and 50 mg/kg Fosfomycin. Since COE-D8 is dissolved in DMSO and Fosfomycin is dissolved in PBS, to exclude the solvent effects, additional groups were included: DMSO group, PBS group, and DMSO + PBS group. After 168 hours, blood samples were collected from the ocular orbit. All blood samples were kept at 4°C overnight, and then centrifuged at 6°C for 15 minutes at 3000 rpm the following day. The serum was used to measure ALT, AST, and BUN levels. These measurements were performed using an automatic biochemical analyzer (Wuhan Servicebio Technology Co., Ltd.). The reference ranges for the three markers were provided by the company.

#### 2.3.7 Embedding tissue sections in paraffin

Euthanize mice by CO_2_ asphyxiation and remove the left kidney, right kidney, liver, and spleen. Place them in 4 mL of 4% paraformaldehyde solution overnight at 4°C. The next day, dehydrate the tissues sequentially in 75% ethanol, 85% ethanol, and 95% ethanol, followed by two rounds of dehydration in 100% ethanol, each for 1 hour. Then, immerse the tissues in xylene for two 1-hour treatments. Finally, immerse the tissues in paraffin wax twice, each for 1 hour, and then embed the tissues in paraffin wax. Subsequently, cut the tissues into 5 μm paraffin sections and mount the sections on glass slides.

#### 2.3.8 Hematoxylin and eosin (H&E) staining

Bake the paraffin sections at 65°C for 2 hours. De-wax the sections twice in deparaffinizing solution, each time for 10 minutes. Wash the sections in 100%, 95%, and 75% ethanol for 10 minutes each, twice in 100%, once in 95%, and once in 75%. Subsequently, stain the sections sequentially with hematoxylin, immerse them in differentiation solution and bluing reagent, and rinse them with running water three times. Then, stain the sections with eosin for 1 minute, followed by two immersions in 95% and 100% ethanol. Finally, incubate the sections in a clearing agent for 4 minutes before mounting with a xylene-based mounting medium.

#### 2.3.9 Bioinformatics

Genomes were extracted from isolated E.coli strains and paired-end 150 bp sequencing using NovaSeq 6000 (Major Biotechnology, Shanghai) was applied. Raw sequencing results were quality controlled using fastp (v0.23.2), where parameters were adjusted according to read length and adapter length. Including read pairs where only one direction passed the filter, the one direction pass quality filter was also included in the assembly construction using SPAdes (v3.15.5). The downloaded assemblies were primarily quality checked using QUAST (v5.2.0)^37^ to filter those with N50 < 200,000 bp.

All assemblies, together with 46 reference genomes ^38^were re-annotated using Prokka (v1.14.6) under fast mode which only detect genes without function annotation. The annotated DNA sequences including coding region, tRNA and non-coding RNA sequences were further clustered by customized pipeline OrthoSLC (v0.2.4).

Meanwhile, this orthologous gene clustering facilitated the comparison of strain SNPs before and after resistance evolution experiment.

Multi-sequence alignment on the clustered core genes were performed using Kalign (v3.3.5) default mode and aligned clusters were later concatenated into core genome of each strain using customized script. The phylogeny computation utilized RAxML (v8.0.2) under GTRCAT mode. 1000 rapid bootstraps were performed on the phylogenetic tree generation.

The structure of genes with amino acid change before and after resistance evolution experiment were predicted by Alphafold2. The molecular docking of COE-D8 and these predicted structures were further simulated using Autodock Vina (v1.5.7) with “exhaustiveness=20”. The docking results were visualized using Pymol(v3.0.5). Data visualization in this study was facilitated by R ggplot2 (v3.4.3), ggtree (v3.10.0) packages.

## 3. Result

### 3.1 Comparison of the Antibacterial Activity of Clinical Antibiotics and COE-D8 against *Uropathogenic Escherichia coli* (UPEC)

A total of 93 *Uropathogenic Escherichia coli* (UPEC) isolates were randomly collected from the hospital between Mar. 2021 to Aug. 2021. The samples represented individuals across various age groups, ranging from a newborn just 11 days old to an elderly individual aged 93 years.(Fig.1a), indicating possible that there may be no inherent correlation between urinary tract infections and age. These 93 strains were tested for their susceptibility to COE-D8 and a variety of clinical antibiotics, including four commonly used first-line antibiotics for treating urinary tract infections (UTIs): Nitrofurantoin, Fosfomycin, Cephalothin, and Compound Sulfamethoxazole. Additionally, three other antibiotics frequently used in the market for managing UTIs—Ofloxacin, Kanamycin, and Ampicillin—were also included in the study(Table S1). All UPEC isolates were resistant to at least two of the antibiotics tested(Fig.1b).

**Figure 1.**
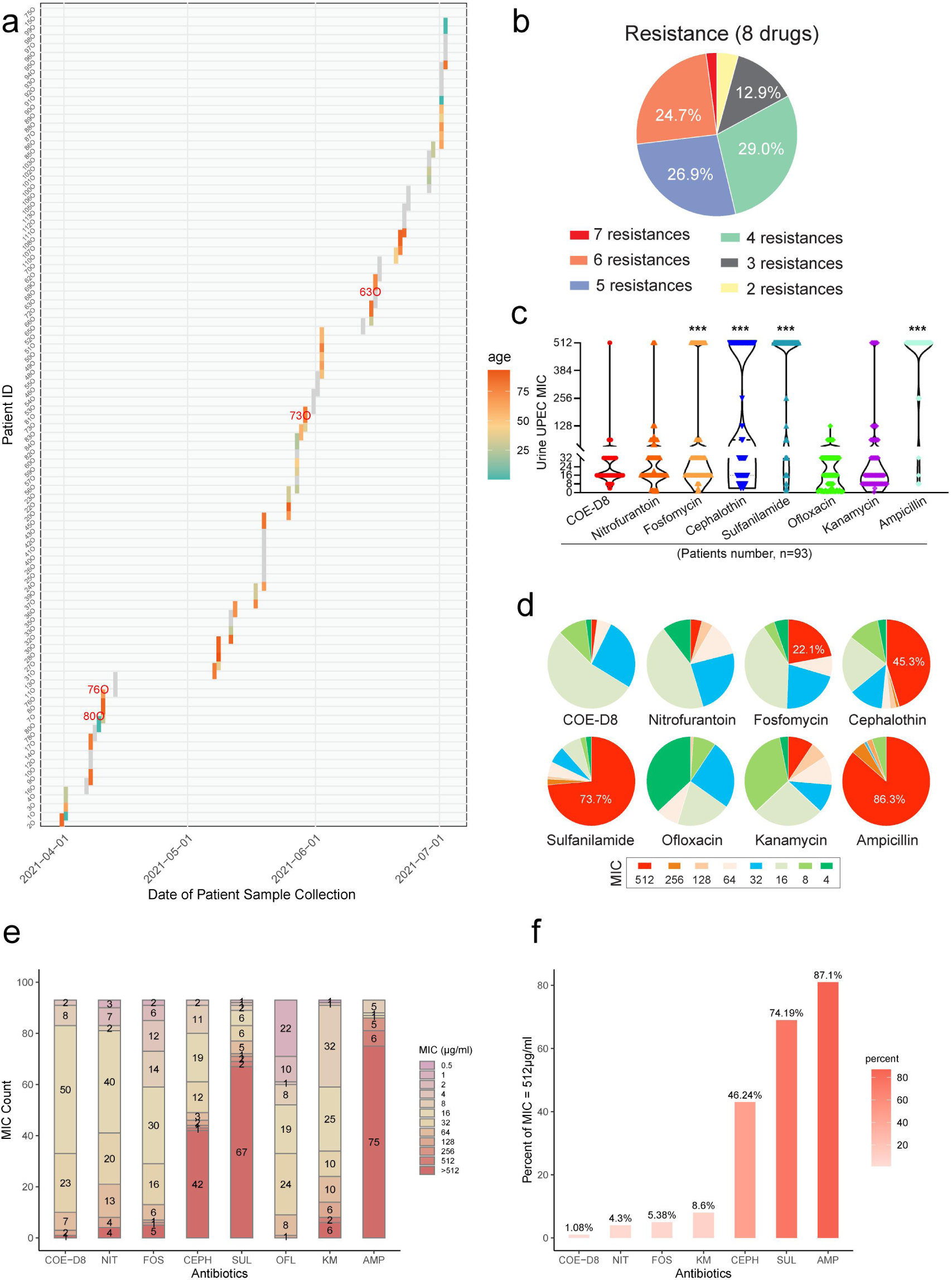
Characteristics of UPEC isolates(n=93) from the Chinese hospital and comparison with COE-D8 antibacterial activity. **a.** Gantt plot showing the duration of samples collection in the urology department of hospital in southwest China. Patient ID are presented on the y axis and the gender of each patient is represented with coloured bars. **b.** Pie chart shows the proportion of resistance counts of each strain. **c.** Violin plot shows the distribution of MICs of COE-D8, Nitrifurantoin, Fosfomycin, Cephalothin, Compound Sulfamethoxazole, Ofloxacin, Kanamycin and Ampicilin for the UPEC isolates. **d.** Eight pie charts show that the MICs concentration proportion of UPEC against COE-D8 and seven other antibiotics. **e.** Stacked chart shows the count of MIC values corresponding to each drug. **f.** Bar chart shows the proportion of antibacterial activity at high drug concentrations for each type of drug against the isolated bacteria.

The isolates had a MIC enrichment region for COE-D8 of 16 μg/mL (wild type MIC= 8 μg/mL), and only one isolate had to be given a high dose (512 μg/mL) to inhibit its growth. The MIC enrichment regions of the remaining seven antibiotics were all far higher than 4 times MIC of the wild type (Fig.1c), indicating that UPEC showed varying degrees of resistance to these seven antibiotics(Fig.1d). However, COE-D8 exhibited significant antimicrobial activity against all 93 UPEC strains, including the highly virulent UTI pathogen UTI89. To be specific, COE-D8 inhibited the growth of 86 UPEC strains at concentrations ranging from 8 to 32 μg/mL, with two strains inhibited at only 4 μg/mL. In contrast, the commonly used antibiotics (Ofloxacin, Kanamycin, and Ampicillin) did not exhibit comparable inhibitory activity against UPEC at the same concentrations. Notably, only 6 out of the 93 UPEC strains (6.5%) required concentrations exceeding 64 μg/mL of COE-D8, with only one strain (1.1%) requiring a concentration above 512 μg/mL. In comparison, the other seven antibiotics failed to produce similar bacteriostatic effects. (Fig.1e).

Furthermore, for the clinically preferred antibiotics—Fosfomycin, Cephalothin, and Compound Sulfamethoxazole—the proportion of strains with a minimum inhibitory concentration (MIC) ≥ 512 μg/mL was alarmingly high: 22.11%, 45.26%, and 73.68% respectively (Fig.1f). This suggests that there is a significant likelihood that patients may not respond to treatment even when high doses of these antibiotics are administered. In contrast, COE-D8 demonstrated superior efficacy, with only one case (1.07%) where the bacterial growth could not be inhibited at a concentration of 512 μg/mL. Moreover, the MIC of COE-D8 was 16 μg/mL for the majority of strains, accounting for 54.8% of the cases—ranking it first compared to the other seven antibiotics. This indicates that, at lower concentrations, COE-D8 could effectively control UPEC infections in most patients.

Therefore, COE-D8 exhibits potent inhibitory effects against UPEC compared to other first-line and commonly used antibiotics.

### 3.2 The phylogeney of UPEC and COE-D8 is anti-UPEC

The experiment described above did not differentiate whether COE-D8 acts to inhibit bacterial growth or to kill bacteria. To address this, we first selected two B2 UPEC strains (#73 and #76), one B1 strain (#63), and one E strain (#80) based on the previous study^38^, while also considering the corresponding MIC values (Table S1). We then inoculated the four Gram-negative UPEC pathogens into growth media and exposed them to 4×MIC concentrations of COE-D8, monitoring bacterial viability over time (Fig. 2). We found that, after 3 hours of treatment with COE-D8, the bacterial count for B2 UPEC strains was reduced to zero. For B1 and UPEC strains, a significant reduction in bacterial count was observed, with effects comparable to those of clinical antibiotics. Based on these experiments, we conclude that COE-D8 exhibits bactericidal activity and is effective against UPEC strains from different evolutionary groups.

**Figure 2.**
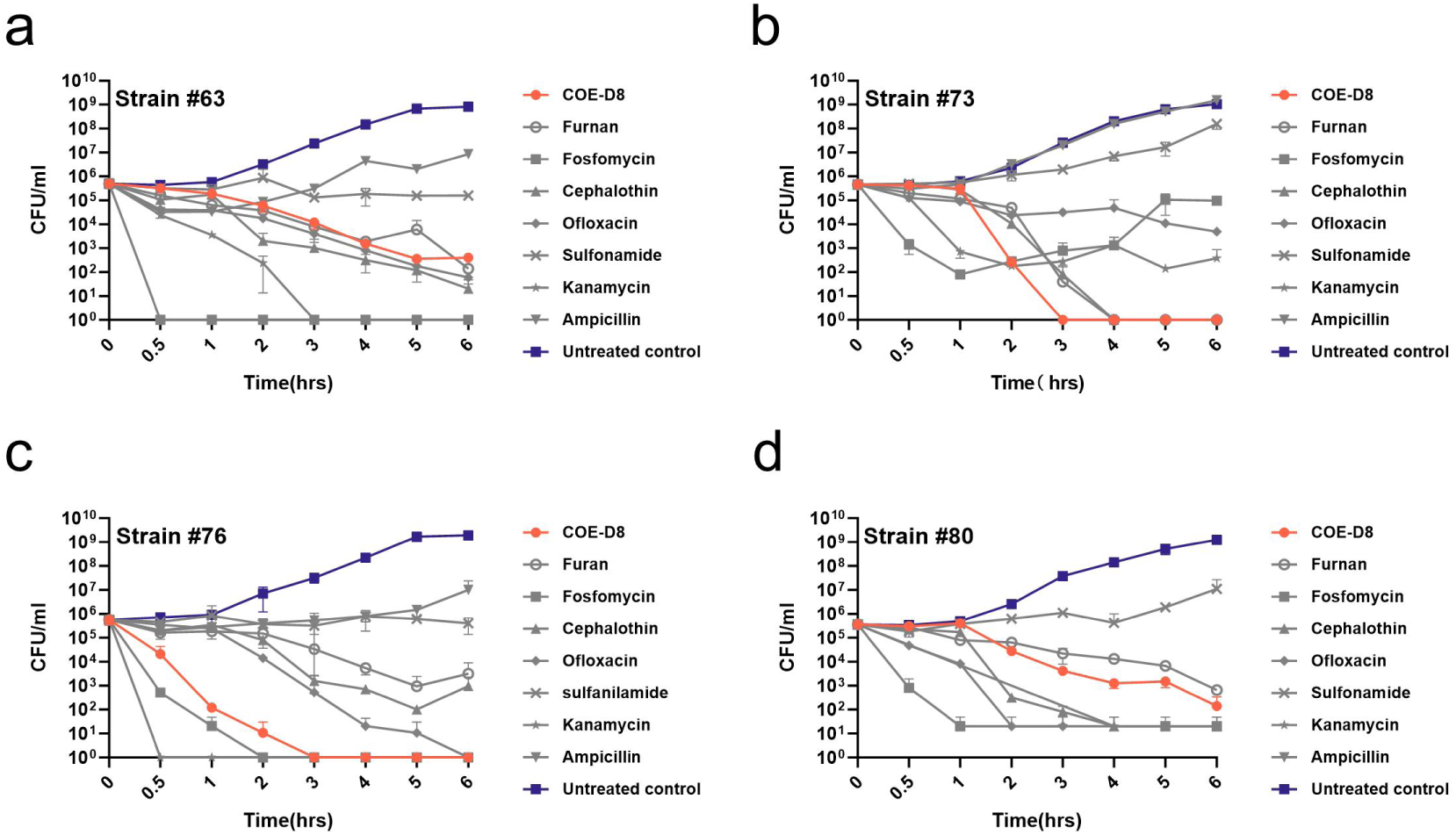
Time-killing assay of 4MIC concentration of COE-D8 and various antibiotic against **a.** Strain#63; **b.** Strain#73; **c.** Strain #76; **d.** Strain #80.

### 3.3 COE-D8 kills bacteria by disrupting the bacterial cell membrane

To further investigate the mechanism of action of COE-D8, we performed repeated passage experiments on selected representative strains. Gradually increased concentrations of COE-D8 were used, and bacteria were isolated from the final passage. After 33 days, the minimum inhibitory concentration (MIC) of COE-D8 for the UPEC strain #76 was found to increase from 1 μg/ml on day 0 to 128 μg/ml (Fig. 3a), indicating that the resistance to COE-D8 increased by 128 times, while the MIC of COE-D8 for strains #63 and #74 was 4 to 16 times higher than their parental counterparts after coresponding 33 days and 19 days (Fig. 3b). Subsequently, whole-genome sequencing was performed on three COE-D8-resistant isolates, which exhibited increased MICs compared to their corresponding non-evolved strains. Single copy core genome phylogeney showed a very close genetic relationship for a experiment pairs (Fig.4). By comparing the total of insertion, deletion, fragment loss/gain and SNPs of strains with their resistant isolate, strain #63 showed 561 bp difference over 14 genes, strain #74 showed 432 bp difference over 15 genes, and strain #76 showed 467 bp difference over 27 genes (Table S2).

**Figure 3.**
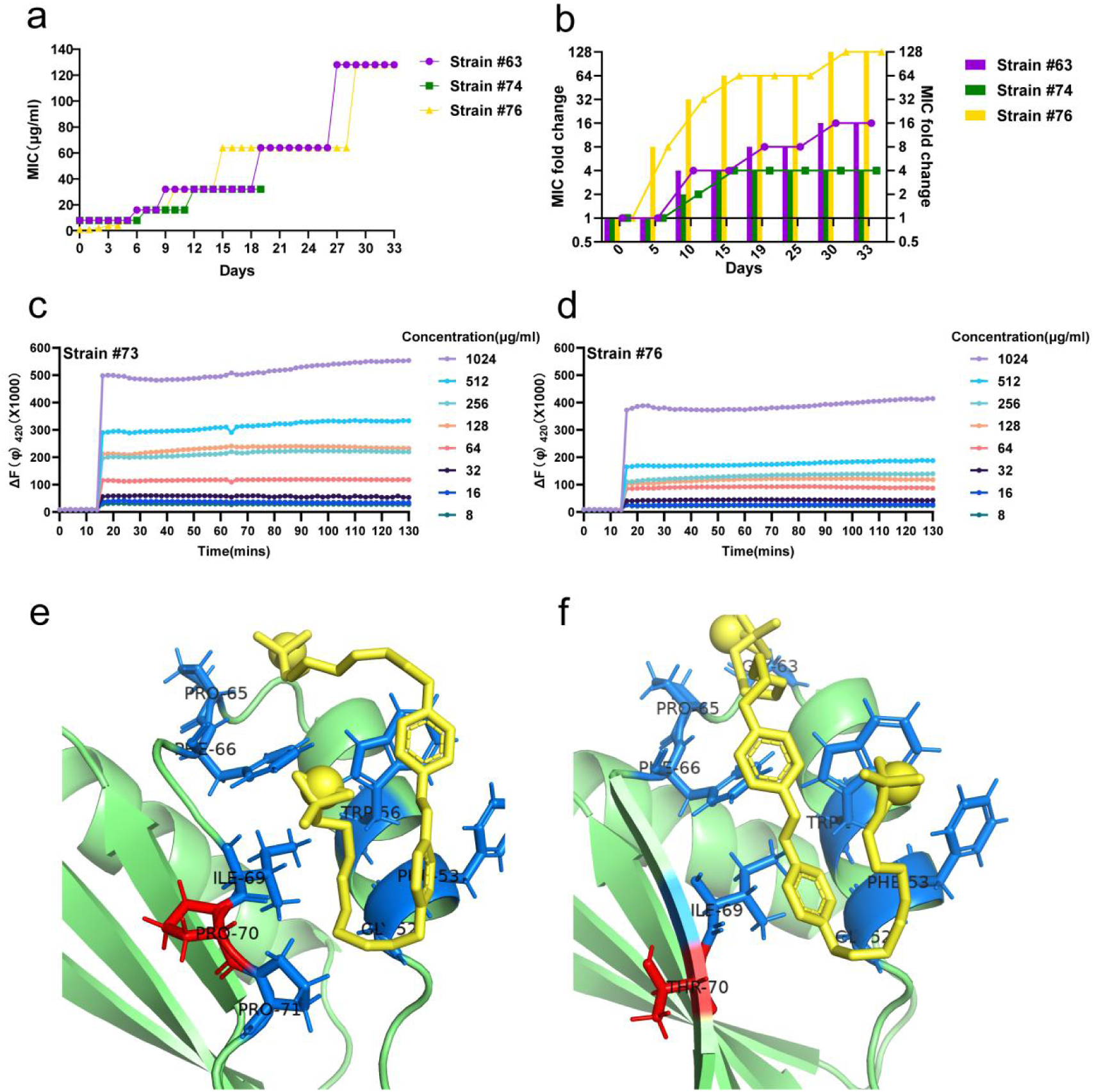
**a. b.** Evolution of COE-D8 resistance with Strain #63 (purple); Strain #74 (green); Strain #76 (yellow). a. line chart shows the MIC change of bacteria. b. the combination of line and bar chart shows the MIC fold change of bacteria: strains are visible growth at the highest concentration of COE-D8 were transferred daily. Data are reported as the highest COE-D8 concentration at which growth was observed and given as the MIC or MIC fold increase in concentration relative to the MIC_90_ on day 1. **c. d.** Relative level of cell membrane electric potential (ΔΨ) of Strain #73(**c**) and Strain #76(**d**) cells exposed to increasing concentrations of COE-D8. **e. f.** are simulated docking results between COE-D8 and *MzrA* of strain #64. While (**e**) is the protein before resistance evolution and (**f**) is the protein after the evolution. The mutated 70^th^ position from proline to theronine was marked as red, the pocket amino acids were marked as blue, and COE-D8 was marked as yellow.

**Figure 4.**
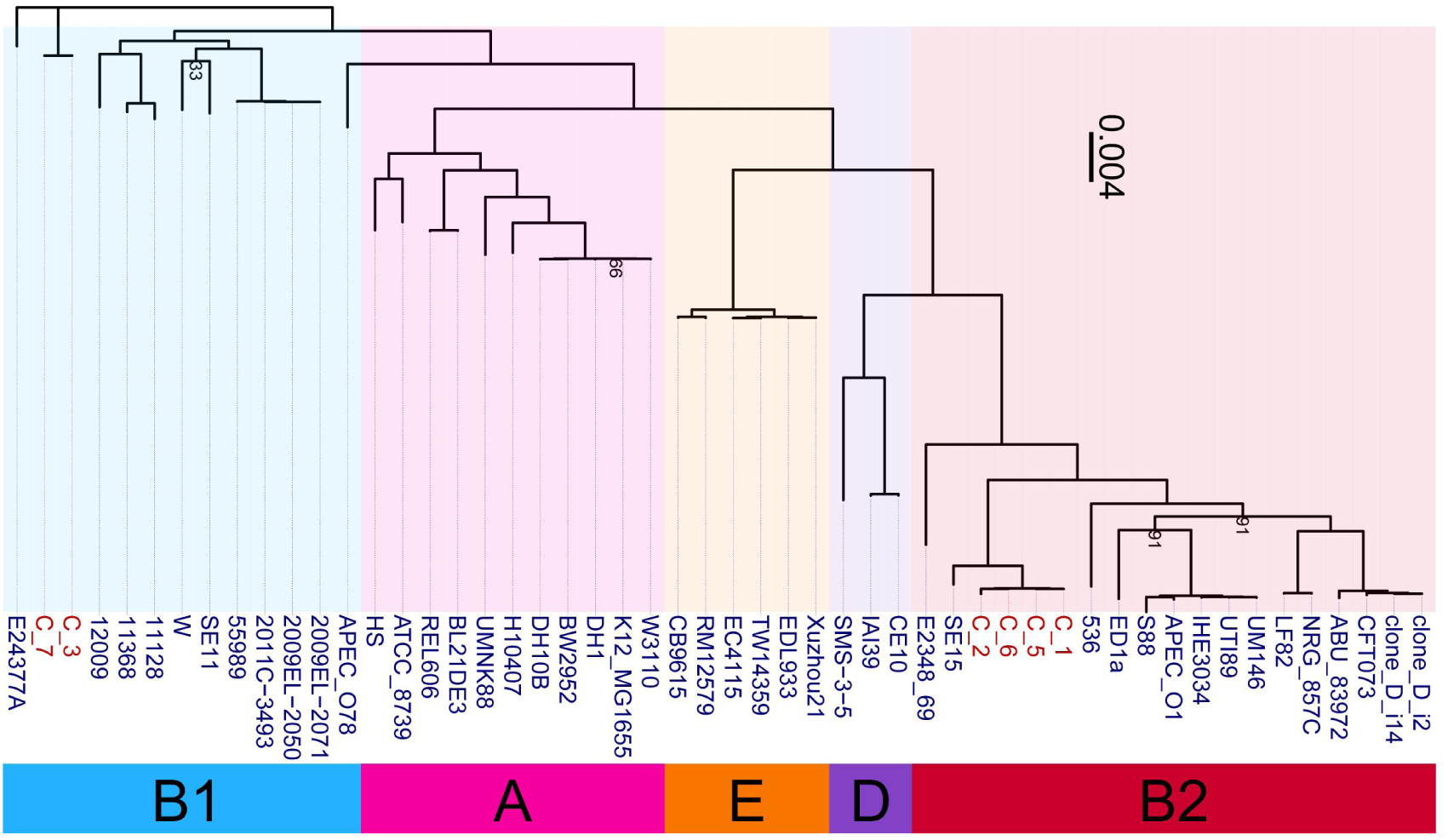
Single-copy core genome based phylogeny of 46 reference (blue strain name) and 3 isolated with there corresponding resistant isolate (red name) E. coli strains. The clade of each cgST was stained differently and corresponding cgST names are in the color bar below. Bootstrap values lower than 95 are showed.

To assess the impact of COE-D8 on outer membrane permeability, we conducted 1-N-phenyl naphthylamine (NPN) uptake fluorescence assays. NPN is a neutral hydrophobic fluorescent probe that is typically excluded by the outer membrane; however, an increase in fluorescence intensity occurs when NPN enters the outer membrane, indicating enhanced membrane permeability^39^. The results showed that COE-D8 induced a dose-dependent increase in fluorescence intensity in all tested UPEC strains, including UPEC #76 (Fig. 3c) and UPEC #63 (Fig. 3d), suggesting that the outer membrane permeability was increased. Thus, COE-D8 kills UPEC by acting on the bacterial cell membrane.

To further explore how those mutated genes could potentially affect the permeability, their protein structure before and after resistance experiment were predicted by AlphaFold2. Cell membrane and cell wall related genes which mutated before and after resistance experiment were evaluated for their affinity with COE-D8 using Autodock Vina. Gene *MzrA* from strain #64, showed a proline to theronine change at its 70^th^ position and changed the neighboring structure from a turn into an β-strand. Over simulated docking attempts by searching the full protein structure, the aromatic rings of COE-D8 could be reproducibly placed next to the β-strand at 70^th^ theronine which was reproducibly not achievable when it is proline at 70^th^. This difference also resulted in a lower docking energy of −5.0 kcal/mol than docking with the protein before resistance evolution −4.6 kcal/mol (Table S3). This could be the target protein specifically affected by COE-D8, and undermining the ability for *MzrA* to regulate peptidoglycan in cell wall synthesis^40^.

### 3.4 Biocompatibility of COE-D8 and combination with Fosfomycin activity

In addition to its antimicrobial properties, biocompatibility with mammalian cells is also a critical factor for antibiotic agents. The *in vitro* cytotoxicity of COE-D8 and a range of antibiotics was assessed using the 3T3 cell line. After a 24-hour treatment, the half-maximal inhibitory concentration (IC_50_) of the antibiotics Kanamycin, Fosfomycin, Ampicillin, Cephalothin, and Ofloxacin was found to be greater than or equal to 512 μg/ml, indicating that the toxicity to mammalian cells is negligible (Fig.5a). Therefore, these antibiotics provide a broad concentration window and can be used as adjuncts without inducing additional toxicity. The corresponding IC_50_ value of COE-D8 was 6 μg/ml. The cytotoxicity changes caused by the combination of COE-D8 with antibiotics were subsequently examined. Supplementation with 512 μg/ml Fosfomycin did not induce any additional toxicity, and after the combination, the IC_50_ of COE-D8 increased to above 16 μg/ml, as the cytotoxicity curve shifted upward (Fig.5b). Thus, based on the IC_50_ in the 3T3 cell line, the IC_50_ increased by approximately threefold (from 6 to 16). These results are consistent with individual cytotoxicity experiments, which showed that 512 μg/ml Fosfomycin alone had almost no toxic effect on 3T3 cells, with cell viability remaining around 100%. Notably, a slightly greater than 100% survival rate was calculated for Kanamycin-treated 3T3 cells in Fig.5a, suggesting that the metabolic activity of the cells may have been enhanced by low concentrations of Kanamycin. In summary, the combined use of COE-D8 with Fosfomycin can achieve improved biocompatibility.

**Figure 5.**
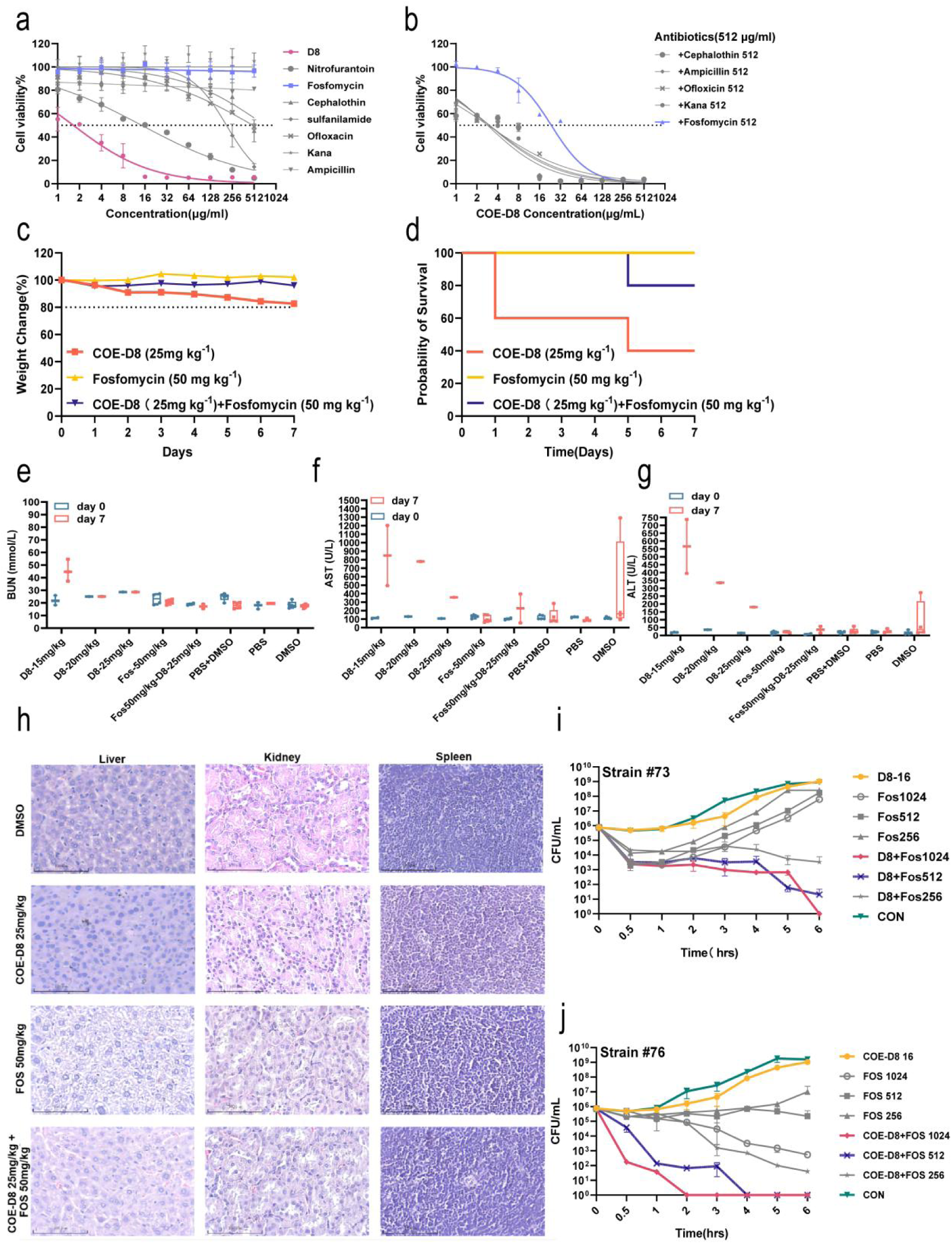
(**a-g**) Cytotoxic effects of COE-D8 and combination with antibiotics *in vivo* and *in vitro*: **a.** *In vitro* cytotoxic efficacy of COE-D8 with 3T3 cell; **b.** *In vitro* cytotoxic efficacy of COE-D8 plus antibiotics (512 μg/ml) with 3T3 cell; **c.** Weight change and **d.** survival% of mice after intraperitoneal injection of COE-D8(25 mg.kg^− 1^), Fosfomycin(50 mg.kg^−1^) and combination of both drugs; Mice viability was followed up to 7 days postinfection; **e.** urea nitrogen (BUN); **f.** aspartate amino transferase (AST); **g.** alanine aminotransferase (ALT) levels in blood from mice injected with COE-D8(15 mg.kg^−1^), COE-D8(20 mg.kg^−1^), COE-D8(25 mg.kg^−1^), Fosfomycin(50 mg.kg^−1^) and combination of Fosfomycin(50 mg.kg^−1^) and COE-D8(25 mg.kg^−1^). PBS and DMSO were used as solvent controls. Blood for drugs-injected mice was drawn at post-injection 0 d and 7 d after administration of the first injection. There were five mice in each group, and data for individual mice were shown as well as means and SDs. **h.** Histological images of main organs of mice at 7 days after drugs (COE-D8 and Fosfomycin) injection: liver, kidney, and spleen. Scale bar=200 μm. Time-killing assay to evaluate antibacterial efficiency of the combination of COE-D8 (16 μg/ml) and Fosfomycin (1024/512/256 μg/ml) against **i.** Strain#73; **j.** Strain #76.

Given the differences between *in vivo* and *in vitro* environments, particularly the potential for combined compounds to induce excessive toxicity effects *in vivo*, the acute toxicity of COE-D8 was evaluated in mice. COE-D8 was administered at a single dose of 25 mg.kg^−1^, Fosfomycin at 50 mg.kg^−1^, and their combination via intraperitoneal injection to three groups of mice. Over a period of 7 days post-injection, no significant changes in body weight were observed in any of the groups (Fig.5c). Mice treated with Fosfomycin and the combination of compounds tolerated the treatment well, with survival rates significantly higher than those observed in the COE-D8-only group (Fig.5d). Furthermore, following the combination treatment, nephrotoxicity was absent (Fig.5e), and hepatotoxicity was reduced by approximately twofold (Fig.5f, g), with reference standards provided by Wuhan Servicebio Technology Co., Ltd, China. These findings confirm that the combination of COE-D8 and Fosfomycin mitigates the acute toxicity of COE-D8 when used alone *in vivo* (Table S4). Histological assessments of major organs from treated animals, including the heart, liver, kidneys, lungs, and spleen, revealed no significant pathological abnormalities or lesions (Fig.5h).

Finally, considering the changes in the bactericidal activity of COE-D8 after the “detoxifying” effect of Fosfomycin, time-point-dependent colony counting experiments were conducted with B2-type UPEC. The results showed that after the addition of a high dose (512 μg/ml) of Fosfomycin, COE-D8 exhibited enhanced bactericidal efficacy against UPEC strains #73 (Fig.5I) and #76 at 1× MIC (Fig. 5J). Notably, Fosfomycin itself did not show bactericidal activity at a concentration of 512 μg/ml. Therefore, the combination of Fosfomycin and COE-D8 not only improves the biocompatibility of COE-D8 but also enhances its bactericidal efficacy.

## 4. Discussion

The increasing severity of antibiotic resistance remains one of the greatest challenges in the treatment of urinary tract infections (UTIs), leading to a rise in the number of recurrent UTI patients. Antibiotic-tolerant bacteria are associated with prolonged treatment durations and recurrent infections. Although some preventive measures have been proposed for recurrent UTIs, their effectiveness remains limited. Therefore, there is an urgent need to develop new antimicrobial compounds to address the growing issue of antibiotic resistance. Surprisingly, COE-D8 was able to inhibit the growth of clinical UPEC strains at a concentration of 16 µg/mL, while Fosfomycin, commonly used in hospitals, could not suppress the growth of clinical UPEC strains even at a high dose of 512 µg/mL. This is not an isolated observation, as it was found in 5.3% of a cohort of 93 patients. Compared to other con antibiotics against UPEC, COE-D8 also demonstrated superior antimicrobial and bactericidal activity. This is likely due to the mismatch in size between COE-D8 and the lipid bilayer, which may result in effective membrane disruption. This finding offers new perspectives for addressing the multi-drug resistance of UPEC. In *in vivo* mouse models, COE-D8, when used in combination with Fosfomycin, reduced liver and kidney damage. However, the treatment still caused a 20% mortality rate in mice, indicating the need for further chemical modifications to reduce toxicity and enhance efficacy.

In the resistance induction study of COE-D8, strain #80 of UPEC, after 28 days of evolution, died, and the strain, which exhibited a 32-fold increase in MIC by day 23, failed to recover. This may be due to the phenomenon of pseudoresistance in strain #80 or to a regulatory mechanism of resistance genes that are expressed only in the presence of COE-D8 but remain dormant otherwise. This hypothesis is supported by the failure of strain #80 to grow in the absence of COE-D8 on the final day. Similarly, strain #80, which was frozen mid-experiment, failed to revive for the same reason. However, the mechanism of drug dependency of resistance genes remains unclear. Subsequent bioinformatic analysis of sequencing data revealed some proteins changed before and after resistance evolution, such as NlpD putative outer membrane lipoprotein, MalT transcriptional activator andmultidrug efflux transporter EmrE, etc., but by docking the protein with COE-D8, it was found that the mutation point did not appear in the docking pocket. The potential reasons that we could not find more direct link between genetic change and resistance increase are: second-generation sequencing is difficult to perfectly restore the genome, and many drug resistance changes may be copy number changes rather than real SNP changes, so the copy number cannot be successfully analyzed in the results; also, because of convergence of evolution^41^, we could not observe a reproducible genetic change that E.coli applied to adapt resistance. In addition, this may be a systematic change during the resistance evolution. The current data is difficult to prove that the mutation site is the best point for drug target bacteria based on mutations from only one or several SNPs.

This work highlights the need for continued exploration of novel antimicrobial agents and the complex mechanisms underlying bacterial resistance, particularly in the context of recurrent UTIs caused by UPEC.

## 5. Conclusion

In conclusion, continued research on the effects of COE-D8 against Gram-negative bacteria, specifically UPEC from clinical patients in Southwest China, demonstrates that COE-D8 exhibits superior antibacterial and bactericidal activity compared to frontline antibiotics (Nitrofurantoin, Fosfomycin, Cephalothin, and Compound Sulfamethoxazole), as well as commonly used antibiotics like Ofloxacin, Kanamycin, and Ampicillin. Furthermore, the combination of COE-D8 with Fosfomycin significantly reduces cytotoxicity both *in vitro* and *in vivo* without compromising the antibacterial efficacy of COE-D8. Whole genome sequencing followed by computational analysis reveals that the potent mechanism of COE-D8 against UPEC is associated with the cell wall protein, which regulates the peptidoglycan in cell wall synthesis, MzrA. It may affect bacterial resistance to COE-D8 by regulating the pathway of cell wall synthesis. When bacteria encounter external antibiotic pressure, the function of MzrA may help maintain the integrity of the cell wall, thereby resisting drug attack. This study represents the first report of COEs in combination with Fosfomycin reducing cellular toxicity to mammalian cells without diminishing bactericidal effectiveness.

## Supporting information

Scheme S1; Table S1;Table S2;Table S3;Table S4

## Acknowledgements

This work was supported by the National Natural Science Foundation of China No.32070818 and 82460142 to C.M.G., Natural Science Foundation of Yunnan Province of China [202001BB050005], the Xingdian talent support program of Yunnan Province to C.M.G., and Guangdong Hybribio Biotech Funding No.H20230314, H20230311 and H20230313 to C.M.G.

## Contributions

C.M.G supervised and guided the overall research; X.Z.X supervised and provided computing power; L.R and C.M.G conceived the project; L.R. present the idea of the whole paper; L.R., C.F. conducted the in vitro and in vivo biological tests; C.F. conducted most of the experiments; J.J.C conducted the sequence analysis, compound docking binding site and modeling analysis; Y.X.W., Y.J.R. and W.L.Y. collected clinic samples and prepared relevant ethics; L.R. and J.J.C drafted the manuscript.

## References

1. Czajkowski, K., Broś-Konopielko, M. & Teliga-Czajkowska, J. Urinary tract infection in women. Przeglad Menopauzalny Menopause Rev. 20, 40–47 (2021).

2. Foxman, B. Epidemiology of urinary tract infections: Incidence, morbidity, and economic costs. Dis. Mon. 49, 53–70 (2003).

3. Foxman, B. The epidemiology of urinary tract infection. Nat. Rev. Urol. 7, 653–660 (2010).

4. Kim, W.-J., Shea, A. E., Kim, J.-H. & Daaka, Y. Uropathogenic Escherichia coli invades bladder epithelial cells by activating kinase networks in host cells. J. Biol. Chem. 293, 16518–16527 (2018).

5. Croxen, M. A. & Finlay, B. B. Molecular mechanisms of Escherichia coli pathogenicity. Nat. Rev. Microbiol. 8, 26–38 (2010).

6. Hybiske, K. & Stephens, R. S. Exit strategies of intracellular pathogens. Nat. Rev. Microbiol. 6, 99–110 (2008).

7. Zowawi, H. M. et al. The emerging threat of multidrug-resistant Gram-negative bacteria in urology. Nat. Rev. Urol. 12, 570–584 (2015).

8. Mancuso, G., Midiri, A., Gerace, E. & Biondo, C. Bacterial Antibiotic Resistance: The Most Critical Pathogens. Pathogens 10, 1310 (2021).

9. Zhang, L., Dhillon, P., Yan, H., Farmer, S. & Hancock, R. E. W. Interactions of Bacterial Cationic Peptide Antibiotics with Outer and Cytoplasmic Membranes of *Pseudomonas aeruginosa*. Antimicrob. Agents Chemother. 44, 3317–3321 (2000).

10. Van Gerven, N., Waksman, G. & Remaut, H. Pili and Flagella. in Progress in Molecular Biology and Translational Science vol. 103 21–72 (Elsevier, 2011).

11. Haraoka, M. et al. Neutrophil Recruitment and Resistance to Urinary Tract Infection. J. Infect. Dis. 180, 1220–1229 (1999).

12. Blanco, M. et al. Detection of pap, sfa and afa adhesin-encoding operons in uropathogenic Escherichia coli strains: Relationship with expression of adhesins and production of toxins. Res. Microbiol. 148, 745–755 (1997).

13. Wijma, R. A., Koch, B. C. P., Van Gelder, T. & Mouton, J. W. High interindividual variability in urinary fosfomycin concentrations in healthy female volunteers. Clin. Microbiol. Infect. 24, 528–532 (2018).

14. Wenzler, E., Ellis-Grosse, E. J. & Rodvold, K. A. Pharmacokinetics, Safety, and Tolerability of Single-Dose Intravenous (ZTI-01) and Oral Fosfomycin in Healthy Volunteers. Antimicrob. Agents Chemother. 61, e00775–17 (2017).

15. Wenzler, E. et al. Phase I Study To Evaluate the Pharmacokinetics, Safety, and Tolerability of Two Dosing Regimens of Oral Fosfomycin Tromethamine in Healthy Adult Participants. Antimicrob. Agents Chemother. 62, e00464–18 (2018).

16. Rouf Khawaja, A., et al. Fosfomycin tromethamine. Antibiotic of choice in the female patient: A multicenter study. Cent. Eur. J. Urol. 68, (2015).

17. Giancola, S. E. et al. Assessment of Fosfomycin for Complicated or Multidrug-Resistant Urinary Tract Infections: Patient Characteristics and Outcomes. Chemotherapy 62, 100–104 (2017).

18. Seroy, J. T., Grim, S. A., Reid, G. E., Wellington, T. & Clark, N. M. Treatment of MDR urinary tract infections with oral fosfomycin: a retrospective analysis. J. Antimicrob. Chemother. 71, 2563–2568 (2016).

19. Neuner, E. A., Sekeres, J., Hall, G. S. & Van Duin, D. Experience with Fosfomycin for Treatment of Urinary Tract Infections Due to Multidrug-Resistant Organisms. Antimicrob. Agents Chemother. 56, 5744–5748 (2012).

20. Bermudez, T. A. et al. Raising the alarm: fosfomycin resistance associated with non-susceptible inner colonies imparts no fitness cost to the primary bacterial uropathogen. Antimicrob. Agents Chemother. 68, e00803–23 (2024).

21. Kot, B. Antibiotic Resistance Among Uropathogenic *Escherichia coli*. Pol. J. Microbiol. 68, 403–415 (2019).

22. Mancuso, G., Midiri, A., Gerace, E. & Biondo, C. Bacterial Antibiotic Resistance: The Most Critical Pathogens. Pathogens 10, 1310 (2021).

23. Habibi, A. & Khameneie, M. K. Antibiotic resistance properties of uropathogenic *Escherichia coli* isolated from pregnant women with history of recurrent urinary tract infections. Trop. J. Pharm. Res. 15, 1745 (2016).

24. Smith, P. A. et al. Optimized arylomycins are a new class of Gram-negative antibiotics. Nature 561, 189–194 (2018).

25. Si, Z., et al. A Glycosylated Cationic Block Poly(β-peptide) Reverses Intrinsic Antibiotic Resistance in All ESKAPE Gram-Negative Bacteria. Angew. Chem. Int. Ed. 59, 6819–6826 (2020).

26. Zhang, Q.-Y. et al. Antimicrobial peptides: mechanism of action, activity and clinical potential. Mil. Med. Res. 8, 48 (2021).

27. Bucataru, C. & Ciobanasu, C. Antimicrobial peptides: Opportunities and challenges in overcoming resistance. Microbiol. Res. 286, 127822 (2024).

28. Hancock, R. E. W. & Sahl, H.-G. Antimicrobial and host-defense peptides as new anti-infective therapeutic strategies. Nat. Biotechnol. 24, 1551–1557 (2006).

29. Bahar, A. & Ren, D. Antimicrobial Peptides. Pharmaceuticals 6, 1543–1575 (2013).

30. Bazan, E. L., Ruan, L. & Zhou, C. Improving the antimicrobial efficacy against resistant Staphylococcus aureus by a combined use of conjugated oligoelectrolytes. PLOS ONE 14, e0224816 (2019).

31. Garner, L. E. et al. Modification of the Optoelectronic Properties of Membranes via Insertion of Amphiphilic Phenylenevinylene Oligoelectrolytes. J. Am. Chem. Soc. 132, 10042–10052 (2010).

32. Yan, H. et al. Influence of molecular structure on the antimicrobial function of phenylenevinylene conjugated oligoelectrolytes. Chem. Sci. 7, 5714–5722 (2016).

33. Zhou, C., et al. Informed Molecular Design of Conjugated Oligoelectrolytes To Increase Cell Affinity and Antimicrobial Activity. Angew. Chem. Int. Ed. 57, 8069–8072 (2018).

34. Zhou, C. et al. A Chain-Elongated Oligophenylenevinylene Electrolyte Increases Microbial Membrane Stability. Adv. Mater. 31, 1808021 (2019).

35. Si, Z. et al. Designer co-beta-peptide copolymer selectively targets resistant and biofilm Gram-negative bacteria. Biomaterials 294, 122004 (2023).

36. Ling, L. L. et al. A new antibiotic kills pathogens without detectable resistance. Nature 517, 455–459 (2015).

37. Gurevich, A., Saveliev, V., Vyahhi, N. & Tesler, G. QUAST: quality assessment tool for genome assemblies. Bioinformatics 29, 1072–1075 (2013).

38. Chen, J. et al. Intra-genomic genes-to-genes correlation enables genome representation. Preprint at 10.1101/2024.06.12.598634 (2024).

39. Lv, Y. et al. Antimicrobial Properties and Membrane-Active Mechanism of a Potential α-Helical Antimicrobial Derived from Cathelicidin PMAP-36. PLoS ONE 9, e86364 (2014).

40. Gerken, H., Charlson, E. S., Cicirelli, E. M., Kenney, L. J. & Misra, R. MzrA: a novel modulator of the EnvZ/OmpR two-component regulon. Mol. Microbiol. 72, 1408–1422 (2009).

41. Heyduk, K., Moreno-Villena, J. J., Gilman, I. S., Christin, P.-A. & Edwards, E. J. The genetics of convergent evolution: insights from plant photosynthesis. Nat. Rev. Genet. 20, 485–493 (2019).

