## Supplementary material for "A Novel COE-D8-Fosfomycin Conjugate Effectively Against First-line Antibiotic Resistant *Uropathogenic Escherichia coli*": Scheme S1; Table S1;Table S2;Table S3;Table S4

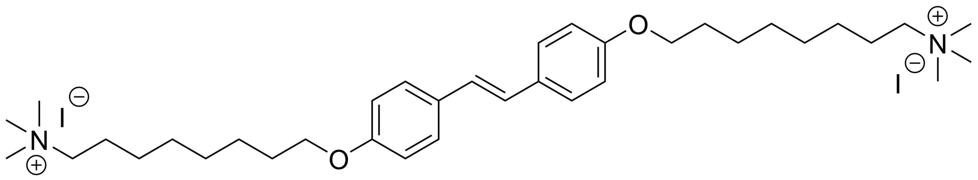

**Scheme S1:** Structure of COE-D8.

| NO. | COE-D8 | NIT | FOS | CEPH | CSUL | OFL | KM | AMP |
|---|---|---|---|---|---|---|---|---|
| **Antimicrobial mechanisms** | Membrane disruption | Cell wall,DNA,RNA synthesis inhibitor | Cell wall synthesis inhibitor | Cell wall synthesis inhibitor | Competitive inhibitor of enzyme dihydropteroate synthetase | DNA gyrase inhibitor | Cell wall synthesis inhibitor | Cell wall synthesis inhibitor |
| **K12** | **8** | 16 | 16 | 16 | >512 | 1 | 4 | 16 |
| **UTI89** | **16** | 16 | 8 | 16 | 16 | 1 | >512 | >512 |
| **YN1** | **32** | 16 | 8 | 32 | 16 | 2 | 8 | >512 |
| **YN2** | **32** | 64 | 16 | 32 | 32 | 2 | 32 | >512 |
| **YN3** | **64** | 64 | >512 | >512 | >512 | 16 | 16 | >512 |
| **YN4** | **32** | 128 | >512 | >512 | 16 | 32 | 64 | >512 |
| **YN7** | **64** | 128 | 8 | 16 | 32 | 2 | 16 | 8 |
| **YN8** | **16** | 64 | 8 | >512 | 16 | 32 | 16 | 512 |
| **YN9** | **32** | 64 | 16 | 16 | 16 | 2 | 16 | 32 |
| **YN10** | **>512** | 64 | 16 | 16 | >512 | 16 | 32 | 512 |
| **YN11** | **16** | 64 | 4 | >512 | 512 | 32 | 64 | 512 |
| **YN12** | **16** | 64 | 16 | 16 | >512 | 1 | 16 | 512 |
| **YN13** | **32** | 16 | 16 | 16 | >512 | 16 | 8 | 256 |
| **YN14** | **16** | 16 | 16 | 16 | >512 | 16 | 8 | 512 |
| **YN15** | **16** | 16 | 16 | 16 | >512 | 1 | 64 | 512 |
| **YN16** | **32** | 16 | 16 | 16 | >512 | 2 | 8 | 8 |
| **YN17** | **16** | 16 | 2 | 16 | >512 | 2 | 16 | 8 |
| **YN20** | **16** | 16 | 8 | >512 | >512 | 2 | 8 | >512 |
| **YN21** | **32** | 16 | 16 | >512 | >512 | 16 | 8 | >512 |
| **YN23** | **8** | 16 | 8 | 16 | >512 | 64 | 8 | >512 |
| **YN24** | **32** | 32 | 64 | 32 | 16 | 64 | 8 | >512 |
| **YN25** | **16** | 16 | 2 | 16 | >512 | 32 | >512 | >512 |
| **YN26** | **32** | 16 | 2 | 64 | >512 | 32 | 0.5 | >512 |
| **YN27** | **16** | 16 | >512 | >512 | >512 | 32 | 8 | >512 |
| **YN28** | **32** | 16 | 16 | 32 | >512 | 32 | 8 | >512 |
| **YN30** | **64** | 16 | 2 | 16 | >512 | 32 | 4 | >512 |
| **YN31** | **32** | 16 | 1 | 16 | >512 | 32 | 8 | >512 |
| **YN32** | **16** | 64 | 8 | 256 | 128 | 64 | 8 | >512 |
| **YN33** | **16** | 16 | 4 | >512 | >512 | 32 | 8 | >512 |
| **YN35** | **16** | 16 | 4 | 32 | 256 | 32 | 8 | >512 |
| **YN36** | **16** | 1 | 1 | >512 | 32 | 32 | 8 | >512 |
| **YN37** | **32** | 2 | 2 | >512 | >512 | 32 | 32 | >512 |
| **YN39** | **32** | 1 | 4 | 512 | >512 | 32 | 16 | >512 |
| **YN40** | **> 64** | 2 | 2 | >512 | >512 | 32 | 64 | >512 |
| **YN41** | **16** | 2 | 4 | >512 | >512 | 32 | 16 | >512 |
| **YN42** | **32** | 2 | 8 | 16 | >512 | 2 | 32 | >512 |
| **YN43** | **16** | 2 | 4 | >512 | >512 | 32 | 16 | >512 |
| **YN45** | **64** | 2 | 256 | 8 | 64 | 16 | >512 | >512 |
| **YN46** | **16** | 2 | 16 | >512 | >512 | 32 | 8 | >512 |
| **YN48** | **16** | 1 | 16 | 8 | >512 | 1 | 16 | >512 |
| **YN49** | **32** | 32 | 64 | >512 | >512 | 32 | 16 | >512 |
| **YN50** | **16** | 16 | 32 | >512 | >512 | 16 | 64 | >512 |
| **YN51** | **32** | 8 | 8 | 8 | 8 | 1 | 16 | >512 |
| **YN52** | **16** | 8 | 32 | 4 | 4 | 8 | 16 | >512 |

| | | | | | | | | |
|---|---|---|---|---|---|---|---|---|
| **YN53** | **32** | 64 | 32 | 32 | 32 | 1 | 16 | >512 |
| **YN54** | **64** | 16 | 8 | >512 | >512 | 1 | 16 | >512 |
| **YN55** | **32** | 64 | 64 | >512 | >512 | 16 | 16 | >512 |
| **YN56** | **16** | 32 | 32 | 8 | 8 | 1 | 32 | 256 |
| **YN57** | **16** | 16 | 32 | 32 | 32 | 1 | 8 | 256 |
| **YN59** | **16** | 64 | 16 | 16 | 16 | 1 | 8 | 256 |
| **YN60** | **16** | 32 | 4 | >512 | >512 | 64 | 32 | >512 |
| **YN62** | **32** | 16 | 4 | 16 | >512 | 1 | 16 | >512 |
| **YN63** | **32** | 16 | 32 | 8 | 512 | 16 | 32 | >512 |
| **YN65** | **> 64** | 16 | 8 | >512 | 64 | 1 | 8 | >512 |
| **YN66** | **16** | 16 | 4 | >512 | >512 | 1 | 16 | >512 |
| **YN68** | **32** | 64 | 16 | >512 | 64 | 64 | 8 | >512 |
| **YN69** | **16** | 16 | 4 | 8 | >512 | 8 | 8 | >512 |
| **YN70** | **64** | 16 | 4 | >512 | 2 | 8 | 16 | >512 |
| **YN72** | **16** | 16 | 32 | >512 | 64 | 32 | 128 | >512 |
| **YN73** | **16** | 128 | >512 | >512 | >512 | 128 | 512 | >512 |
| **YN74** | **4** | 16 | 16 | >512 | >512 | 16 | 128 | >512 |
| **YN76** | **4** | 16 | 8 | >512 | 256 | 16 | 64 | >512 |
| **YN78** | **16** | 16 | 16 | 32 | >512 | 2 | 128 | >512 |
| **YN80** | **8** | 16 | 16 | 4 | >512 | 16 | >512 | >512 |
| **YN81** | **8** | 64 | 32 | >512 | >512 | 32 | >512 | >512 |
| **YN82** | **16** | 16 | 32 | 32 | >512 | 2 | 128 | >512 |
| **YN83** | **16** | 16 | 16 | >512 | >512 | 32 | 64 | >512 |
| **YN84** | **16** | 16 | 16 | 8 | >512 | 1 | 64 | >512 |
| **YN85** | **8** | 32 | 4 | 16 | >512 | 8 | 8 | >512 |
| **YN86** | **16** | 32 | 32 | 32 | >512 | 1 | 8 | 16 |
| **YN87** | **16** | 32 | 16 | >512 | >512 | 8 | 16 | >512 |
| **YN88** | **16** | 128 | 32 | >512 | >512 | 64 | 16 | >512 |
| **YN89** | **16** | 32 | >512 | >512 | >512 | 32 | 16 | >512 |
| **YN90** | **64** | >512 | 16 | >512 | >512 | 64 | 8 | >512 |
| **YN91** | **16** | >512 | 16 | >512 | >512 | 1 | 8 | >512 |
| **YN92** | **16** | 32 | 8 | >512 | >512 | 16 | 64 | >512 |
| **YN93** | **8** | 16 | 32 | 128 | >512 | 1 | 8 | >512 |
| **YN94** | **16** | 16 | 32 | 16 | >512 | 16 | 8 | 8 |
| **YN95** | **16** | 32 | 64 | >512 | >512 | 64 | 128 | >512 |
| **YN96** | **8** | 32 | 16 | 64 | 64 | 1 | 64 | >512 |
| **YN97** | **16** | 32 | 32 | 16 | >512 | 8 | 32 | >512 |
| **YN98** | **16** | 32 | 64 | 128 | >512 | 4 | 32 | >512 |
| **YN99** | **16** | 16 | 64 | 8 | >512 | 1 | 8 | 8 |
| **YN100** | **16** | 32 | 16 | 32 | >512 | 1 | 8 | >512 |
| **YN101** | **32** | 32 | 32 | 8 | >512 | 1 | 8 | >512 |
| **YN102** | **32** | 16 | 8 | >512 | >512 | 16 | >512 | >512 |
| **YN103** | **16** | 32 | 16 | >512 | >512 | 8 | >512 | >512 |
| **YN105** | **16** | >512 | 16 | >512 | >512 | 16 | 16 | >512 |
| **YN106** | **16** | >512 | 32 | >512 | >512 | 16 | 16 | >512 |
| **YN107** | **8** | 32 | 8 | 32 | >512 | 8 | 512 | >512 |
| **YN108** | **8** | 16 | 16 | 64 | 32 | 1 | 32 | >512 |
| **YN110** | **16** | 32 | 16 | >512 | >512 | 16 | 128 | >512 |
| **YN111** | **16** | 16 | 128 | >512 | >512 | 1 | 8 | >512 |
| **YN112** | **16** | 32 | 16 | 8 | >512 | 16 | 16 | >512 |
| **YN113** | **16** | 32 | 16 | 8 | >512 | 32 | 8 | 256 |

**Table S1.** Antimicrobial activity of COE-D8 and other 7 antibiotics against totally 93 UPEC and 2 reference strains( K12 and UTI89). NIT: Nitrofurantoin; FOS: Fosfomycin; CEPH: Cephalothin; CSUL: Compound Sulfamethoxazole; OFL: Ofloxacin; KM: Kanamycin; AMP: Ampicilin.

| Strains | Genes | Mutations during resistance evolution |
|---|---|---|
| Strain 63 | efflux RND transporter permease AcrB | 515W->stop codon |
| Strain 63 | NlpD putative outer membrane lipoprotein | fragment loss from 368 to 380 |
| Strain 63 | MzrA | 70P->T |
| Strain 63 | MalT transcriptional activator | 319C->Y |
| Strain 74 | multidrug efflux transporter EmrE | 3L->P |
| Strain 74 | MalT transcriptional activator | 439S->I |
| Strain 76 | 3,4-dihydroxy-2-butanone 4-phosphate synthase | 100D->E |

**Table S2.** All single base substitutions were non-synonymous mutations coding for either an amino acid substitution. The last column shows the gene before resistance evolution (lefe of right arrow) mutate to the gene on the right hand side experiencing corresponding evolution day.

| Gene | Ligand | Affinity (kcal/mol) |
|---|---|---|
| MzrA[1] | | -5 |
| MzrA[2] | | -4.6 |
| MalT transcriptional activator[1] | | -7.1 |
| MalT transcriptional activator[2] | | -7.2 |
| multidrug efflux transporter EmrE[1] | COE-D8 | -4.2 |
| multidrug efflux transporter EmrE[2] | | -4.1 |
| 3,4-dihydroxy-2-butanone 4-phosphate synthase[1] | | -4.5 |
| 3,4-dihydroxy-2-butanone 4-phosphate synthase[2] | | -4.5 |

**Table S3.** Docking Affinity in Autodock Vina. The gene of MzrA shows a lower docking energy of -5.0 kcal/mol than docking with the protein before resistance evolution -4.6 kcal/mol. There are no docking energy change before and after resistance evolution among the gene of 2003, 3437 and 1752. [1] represent the gene before resistance evolution. [2] represent the gene after resistance evolution.

| | AST (U/L) | | ALT (U/L) | | BUN ( mg/dl ) | |
|---|---|---|---|---|---|---|
| | reference standards (**36.31-235.48 U/L**) | | reference standards (**10.06-96.47 U/L**) | | reference standards (**10.81-34.74mg/dl**) | |
| | **0 day** | **7 day** | **0 day** | **7 day** | **0 day** | **7 day** |
| **COE-D8 25mg /kg** | 108.22 | 356.55 | 15.75 | 180.3 | 22.70 | 28.64 |
| **FOS 50mg /kg** | 121.05 | 79.38 | 17.83 | 23.19 | 23.09 | 20.71 |
| **COE-D8 25mg/kg+FOS 50 mg/kg** | 103.22 | 227.30 | 8.26 | 36.83 | 19.00 | 17.20 |

**Table S4.** *In vivo* safety profile of COE-D8, Fosfomycin and combination of both drugs after intravenous injection.The single or combination drugs do not cause significant acute damage to kidney function, nor do they interfere with the electrolyte balance in the blood at COE-D8 25mg/kg, FOS 50mg/kg and combination injection. Hepatotoxicity was present at COE-D8 25mg /kg-only injection, but the toxicity was reduced by approximately twofold at the combination. Data are expressed as mean standard deviation (n=3). P>0.05 for comparison after 7 day repetitive treatment to 0 day BUN group; statistical analysis was performed using Student's t-test; differences are considered statistically significant with probability P<0.05. ALT, alanine transaminase; AST, aspartate transaminase; BUN, urea nitrogen.
